# Using digital holographic microscopy (DHM) to monitor effects of extracellular matrix (ECM) glycation on cancer cell morphology and migration

**DOI:** 10.64898/2026.07.09.737564

**Authors:** Anupom Deb Nath, Estelle Leclerc, Stefan Vetter

## Abstract

The extracellular matrix (ECM) is a complex network of ubiquitously present acellular material that plays a critical role in cell proliferation, migration, invasion, and tissue morphogenesis. Non-enzymatic glycation of ECM modifies the structure and function of ECM proteins and can support a pro-inflammatory milieu in the tumor microenvironment. However, the impact of glycated ECM on cancer cell growth remains underexplored despite its importance in facilitating disease progression. Here, we investigate the effect of ECM glycation on cancer cell morphology and migration behavior. We used methylglyoxal (MG) as a glycation agent and collagen as our ECM model protein. For in vitro growth analysis, breast cancer cells were seeded on growth surfaces coated with both non-glycated and glycated collagen. Cell behavior was monitored for 24 hours using a real-time holographic imaging system. Holographic image analysis revealed significant differences between non-glycated and glycated growth substrates in cell spreading area, eccentricity, perimeter length, optical thickness, and optical volume, as well as cell migration and motility, which directly influence cell adhesion and proliferation. These changes were found to be cell line biased. Overall, our findings suggest that ECM glycation has a significant effect on cell morphology, migration and cell growth. Holographic live cell imaging was determined to be an excellent method to monitor cells without the need for any labeling and with minimal perturbations.

## Introduction

The extracellular matrix (ECM) is a complex network of ubiquitously present acellular components including collagens, elastin, fibronectin, proteoglycans, and several other glycoproteins that help build structures for the cells and their surroundings^1^. ECM provides biophysical cues to cells, and with different types of receptors such as integrins, cells are able to adhere to ECM^2^. In addition to maintaining cellular homeostasis, ECM provides biochemical signals that play a critical role in cell proliferation, migration, invasion, and tissue morphogenesis^3^. Because ECM molecules are biologically long-lived, their probability of accumulating modifications by glycation or the formation of advanced glycation end products (AGEs) is high^4^. Protein glycation is a time-dependent, non-enzymatic modification of nucleophilic groups within proteins, caused by reactive carbonyl compounds including reducing sugars; it is a complex series of chemical reactions collectively known as the Maillard reaction^5^. AGE formation in the ECM leads to covalent intra- and inter protein crosslinks and modifies the structure and function of ECM in ways that fuels inflammation in the tumor microenvironment, which in turn favors cancer cell proliferation, migration as well as tumor evasion leading to metastases^6^. Tumor-associated ECM modification can occur by ECM deposition, fiber alignment, and crosslinking, leading to increased integrin signaling and favoring tumor cell survival by evading tumor-suppressing mechanisms^7^. Tumor’s metastatic condition not only changes the production of tumor-derived ECM but also the stromal contribution of the tumor ECM by directing the stromal cells to express different sets of ECM proteins^8^. This ECM modification can change its functionality either by modifying its recognition pattern to specific proteins or by altering its mechanical properties, such as causing stiffness to tissues, and there is substantial evidence that suggests changes in ECM functionality play a critical role in tissue inflammation, tumor progression, and metastasis^9,10^.

Non-enzymatic glycation of ECM induces highly invasive cell phenotypes and epithelial-mesenchymal transition (EMT) by promoting transforming growth factor TGF-β^11^. Acerbi et al. demonstrate the stiffening of ECM in mouse models of mammary cancer as well as an increased deposition and thickening of collagen in breast cancer transformation. This study also showed a positive correlation between stroma stiffness and the expression of cellular TGF-β signaling^10^. In addition, glycated ECM exhibits alteration in vascular growth and structural integrity found similar in tumor vasculature structure^12^. In diabetes or hyperglycemic conditions, the rate of ECM glycation are higher, contributing to a source of AGE-modified proteins that subsequently activate the receptor for advanced glycation end products (RAGE)^9^. The impact of glycated ECM on cancer cell growth remains underexplored, considering the extent of ECM glycation and the significant role of AGEs in various diseases and inflammation^13-20^, it is imperative to investigate these effects at the cellular level using high-content approaches.

Collagen is the most abundant fibrous protein found in mammals, comprising around 30% of total protein mass^21^. To date, 28 types of collagens have been identified, and most of them can form a triple-stranded helix form. It is the main structural element of ECM and functions by providing tensile strength, regulating cell adhesion, and maintaining tissue development^22^. Like most ECM components, collagen has a long half-life, making it susceptible to crosslinking and other types of modifications. From an amino acid standpoint, collagen is characterized by high glycine (∼30%) and proline/hydroxyproline content (∼20%), along with elevated amounts of arginine and lysine residue^23^. The arginine and lysine of collagen are most vulnerable to non-enzymatic glycation reactions, resulting in the formation of AGEs. Accumulation of AGEs highly compromises tissue integrity, cell-matrix adhesion, and healing capacity, and also contributes to several human pathologies such as diabetes, arthritis, and cancer^24^.

Digital Holographic Microscopy (DHM) is a label-free imaging technique that allows real-time imaging and continuous morphological evaluation of live cells. The technique involves splitting a laser into two beams-a reference beam and an object beam. The reference beam remains unchanged, while the object beam passes through the specimen. Combining these beams produces an image through the interpretation of the resulting interference pattern, where pixel intensity signifies optical thickness. Optical thickness is influenced by both physical thickness and optical density. Standard image segmentation techniques are employed to identify cells, and various cell features such as optical volume, optical thickness, and perimeter length are extracted. This method allows the capture of quantitative phase images over time, facilitating the tracking of DHM-derived features as cells undergo transformations^25,26^. DHM-derived morphological analysis has been used previously to characterize the state of cells in multiple diseased conditions^27,28^.

In this study, we investigate how the glycation of collagen as a representative ECM protein affects the behavior of cancerous cells. Using continuous, label-free, non-invasive, holographic microscopic imaging of the cells, we are able to characterize cancer cell lines based on phenotypical observations. We used methylglyoxal (MG), a well-known glycolysis by-product, to glycate the most abundant matrix protein, collagen. In-vitro glycation with MG is widely used as a model for in-vivo protein glycation in multiple diseases^29,30^. Here, we presented how glycation affects the ECM protein collagen by amino acid analysis and transmission electron microscopy, then we demonstrated the effect of ECM glycation on non-cancerous and cancerous cells using a holographic imaging system and how it affects cell morphology, migration, and adhesion, finally, we tried to see if the glycated ECM protein has any effect on the gene expression of the cells. Our findings suggest that glycation of ECM proteins significantly alters cancer cells’ morphology and migration patterns that may influence the cancer cells’ growth and metastasis.

## Results

### Amino acid analysis

In this experimental study, collagen with a concentration of 1 mg/ml was subjected to glycation by treatment with 20 mM MG for 3 days. After the glycation process, amino acid analysis was conducted using mass spectrometry to discern alterations in the collagen contents. The findings of the analysis revealed a substantial reduction, amounting to 90% (Fig. 1c), in the signals corresponding to arginine (Arg) residues in the glycated collagen (GC) (Fig. 1b) when compared to the native, non-glycated collagen (NC) (Fig. 1a). This observed reduction in Arg signals suggests a significant impact of glycation on the specific amino acid composition of collagen, thereby highlighting potential modifications. Previous studies reported that MG predominantly modifies the Arg side chain of proteins, resulting in the formation of AGEs^31-33^. Non-enzymatic modification of Arg residues, which is essential for nicotinamide adenine dinucleotide phosphate (NADPH) production, causes disruption in cellular redox balance and has implications on other pathogeneses^34^. Multiple studies reported the MG-mediated modification of Arg residues of hemoglobin associated with physiological complications^35,36^. MG is a naturally occurring byproduct of metabolism found in all cells under both normal and pathological conditions, and it was recognized as the most potent glycation agent, which contributes to structural changes and dysfunction within cells and has implications on diabetes, cardiovascular disease, and cancer^30,34,37-39^. This study reiterates that the collagen we used for morphology analysis was indeed modified due to the glycation process.

**Fig. 1.**
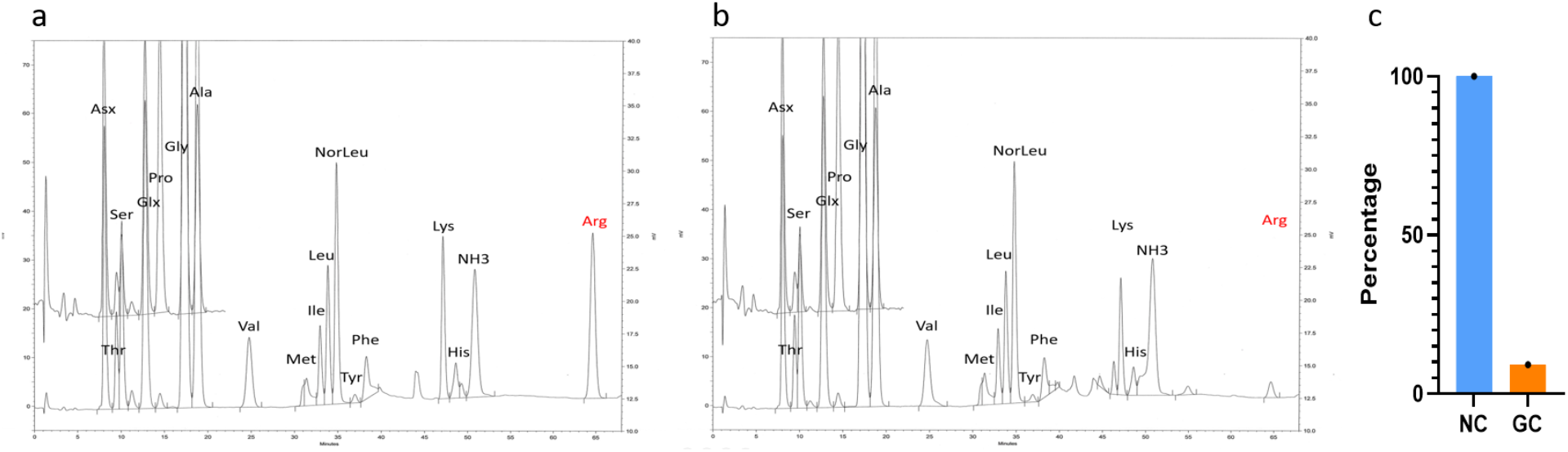
From the amino acid analysis result, the presence of high glycine and proline residues confirms the representative collagen materials. In the GC (Fig. 1b), there was a significant, almost 90% reduction (Fig. 1c) in the absorbance peak for Arg, which was reported to be strongly modified due to non-enzymatic glycation by MG. There was also moderate modification of lysine residues observed.

### Transmission electron microscopy (TEM)

After the validation of amino acid side chain modifications resulting from protein glycation, it became imperative to investigate the consequential structural alterations in the GC. It has been well reported that glycation of ECM protein changes the molecular and structural characteristics of the ECM. Such as glycation of collagen reduces stress tolerance, viscoelasticity, and collagen fiber sliding; these structural changes have significant biological consequences^40,41^. In our experimental study, we wanted to see if glycation of collagen alters the ultrastructure of fibril formation. To assess these changes, NC and GC samples were affixed onto microscopic grids and subjected to analysis using Transmission Electron Microscopy (TEM). This technique allowed for a detailed examination of the ultrastructural features of the collagen specimens at a microscopic level. Upon image analysis, it was discerned that there were no observable structural distinctions between the NC (Fig. 2a) and GC (Fig. 2b) fibrils^42^.

**Fig. 2.**
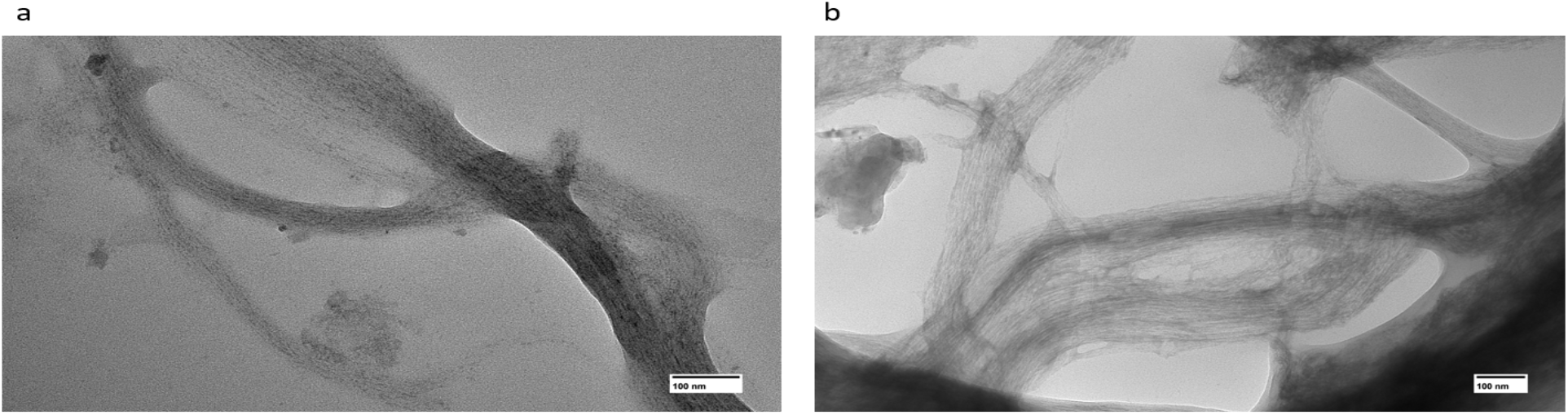
The TEM result demonstrates that the fibril formation capacity of the collagen is not significantly impacted due to the glycation reaction, although glycation impacts the physical properties of collagen in the molecular level. Additional GC TEM images are presented in Supplementary Fig. 1 online.

### Immunoreactivity of the monoclonal anti-methylglyoxal (anti-MG) antibody

MG is the most important AGE precursor and exclusively reacts with the arginine residues of the proteins. Hydroimidazolone (MG-H1) is the major product of MG glycation, resulting from the reaction of MG with arginine.^5,43^ To evaluate the effect of immunoreactivity of MG-mediated ECM glycation, we conducted a non-competitive ELISA assay. In this assay, the anti-MG antibody was utilized to measure the binding affinity with the GC samples. The resultant data, as depicted below (Fig. 3), illustrate a dose-dependent interaction between the anti-MG antibody and the GC. This observed dose-dependent binding pattern provides evidence that the antibody exhibits specificity in recognizing and binding to the GC molecules.

**Fig. 3.**
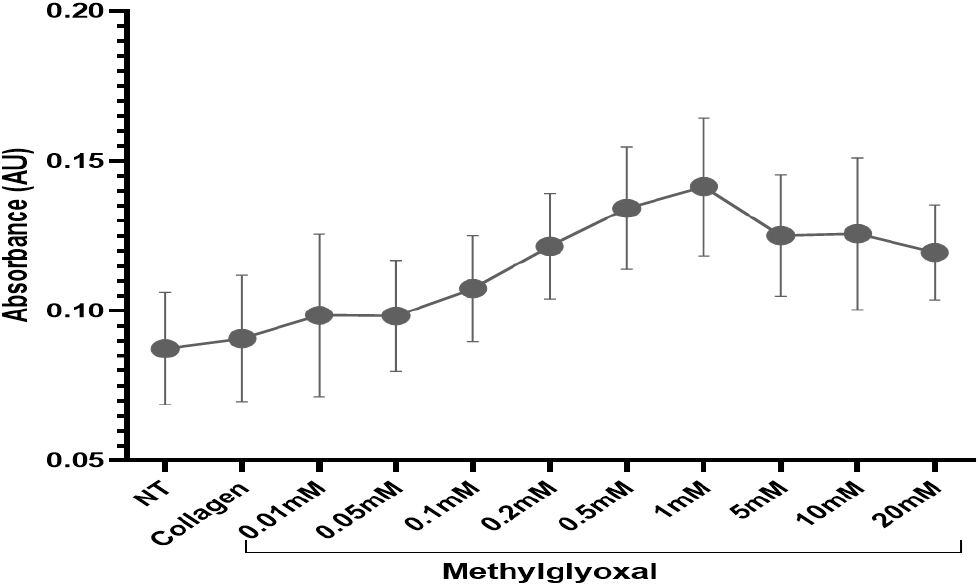
Dose-dependent MG glycation of collagen protein is recognized by anti-MG antibody. Data presented mean absorbance ± SD from N=3 independent experiments.

### Identification of DHM-derived features

To establish the DHM-derived features for cancerous cell behavior, initially, HEK293 and HEK-RGAE cells were cultured on ECM-coated growth culture surfaces. Cells were grown on ECM-coated (Collagen-I, Collagen-IV, Fibronectin, Matrigel) surface along with BSA and a non-treated surface as controls. After 24 hours of post-seeding, cell morphological features were monitored for 24 hours using Holomonitor AppSuite 3.5 software by randomly selecting positions in the culture plate. First, we analyzed the optical thickness of the cultured cells to evaluate comparable differences among the growth surfaces, as optical thickness is a fundamental measurement of DHM. Optical thickness is the measurement of the phase shift when the laser passes through the objects, typically cells. We found comparable differences in optical thickness among the growth surfaces (Fig. 4. data represents the optical thickness of three treatment conditions used for the following cancer cell morphology study) with consistent and reproducible variations.

**Fig. 4.**
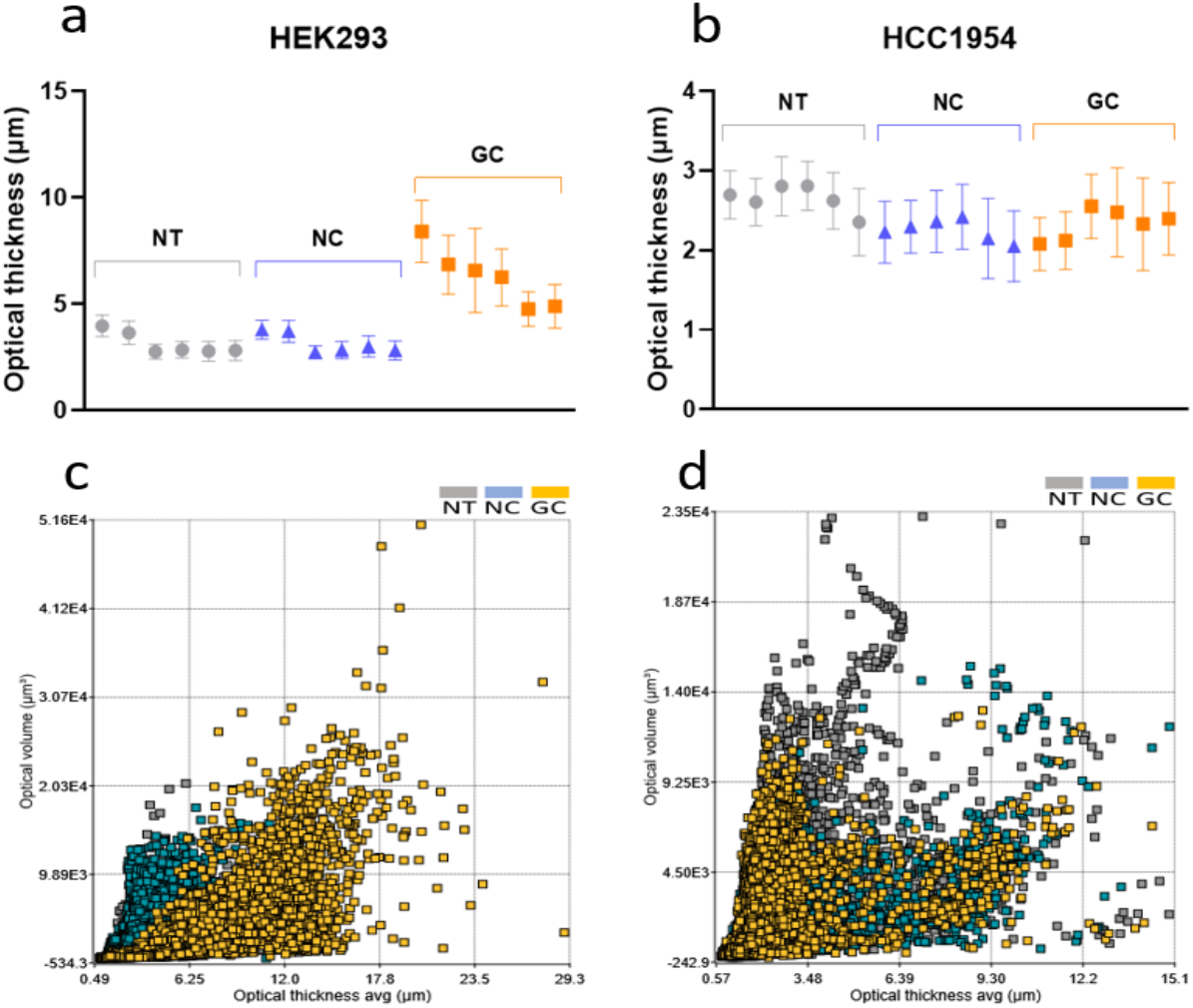
Differences in optical thickness among treatment conditions for HEK293 (Fig. 4a) and HCC1954 (Fig. 4b) cells. Three different growth conditions used for the morphology analysis are presented in the figure. For each cell line, data were presented from 3 independent experiments with replicates. Non-treated (NT) growth surface, non-glycated collagen (NC) coated growth surface, and glycated collagen (GC) coated growth surface. Average optical thickness plotted against optical volume for cells grown on different growth surfaces-HEK293 cells (Fig. 4c) and HCC1954 (Fig. 4d).

**Table 1.** Mean optical thickness (µm) of corresponding Fig. 4 (a, b) for HEK293 and HCC1954 cells grown on NT, NC and GC growth surface.

|  |  | HEK293 |  |  |  | HCC1943 |  |  |
| --- | --- | --- | --- | --- | --- | --- | --- | --- |
|  |  | NT | NC | GC |  | NT | NC | GC |
| EX1 | Rep 1 | 3.97 | 3.80 | 8.40 |  | 2.70 | 2.23 | 2.08 |
|  | Rep 2 | 3.65 | 3.71 | 6.85 |  | 2.60 | 2.30 | 2.12 |
| EX2 | Rep 1 | 2.76 | 2.74 | 6.57 |  | 2.80 | 2.36 | 2.55 |
|  | Rep 2 | 2.85 | 2.83 | 6.25 |  | 2.81 | 2.42 | 2.48 |
| EX3 | Rep 1 | 2.79 | 3.00 | 4.76 |  | 2.62 | 2.15 | 2.33 |
|  | Rep 2 | 2.81 | 2.82 | 4.89 |  | 2.35 | 2.05 | 2.39 |

Both HEK293 and HEK-RAGE (see Supplementary Fig. 3, online) cells grown on ECM-coated surface have significantly reduced optical thickness compared to cells grown in BSA and a non-treated plastic surface. Cells grown in collagen I and fibronectin-coated growth surfaces show significant variations in almost all features compared to control groups. Such as the average cell spreading area for NT-777.7 µm^2^ (Fig. 5a); BSA-780.8 µm^2^; collagen I-946.3 µm^2^ (Fig. 5b); collagen IV-817.4 µm^2^; fibronectin-958.6 µm^2^; Matrigel-812.2 µm^2^. On the other hand, cells grown in collagen IV and Matrigel show significant variations only in optical thickness and optical volume. Such as the average optical thickness for NT-4.7 µm; BSA-5.9 µm; collagen I-2.7 µm; collagen IV-4.0 µm; fibronectin-3.1 µm, and Matrigel-4.5 µm.

**Fig. 5.**
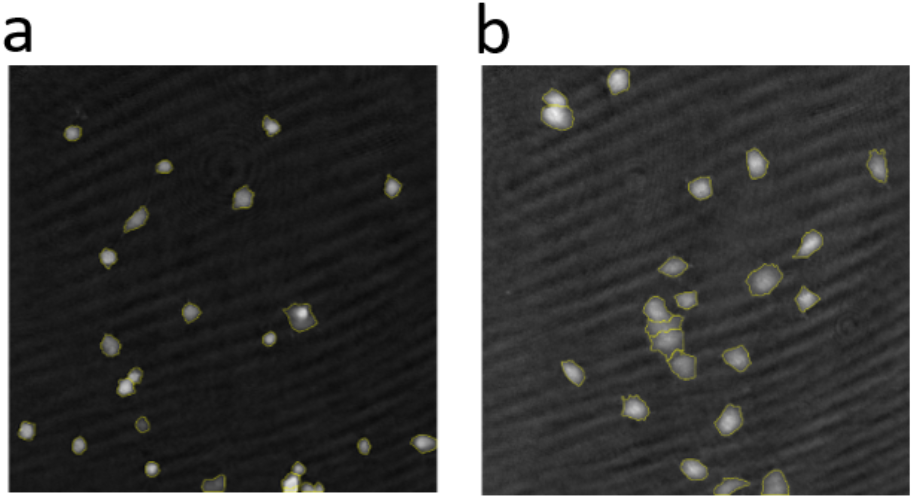
Holographic image of HEK293 cells grown in different growth conditions. Differences in cell spreading area between non-treated growth surface (Fig. 5a) vs cells grown in collagen I treated growth surface (Fig. 5b). Additional cell growth images on different ECM matrices are presented in Supplementary Fig. 2 online.

Once we establish that differences in cell growth on ECM surfaces can be evaluated with reproducibility, we then proceed to check the cell growth on glycated ECM growth surface. For this, HEK293 (see Supplementary Fig. 4, online) cells were grown in three conditions: non-treated (NT) growth surface, NC-coated growth surface, and GC-coated growth surface. Following 24 hours of cell seeding, attached cells were continuously monitored for 24 hours.

Cell morphology and migration pattern analysis revealed there are significant differences in cells grown on NT growth surface vs cells grown on NC-coated growth surface and GC-coated growth surface (see Supplementary Fig. 5, online). Cells grown on GC growth surface have significantly higher optical thickness and optical volume compared to cells grown on NT and NC coated growth surface; NT-3.1µm; NC-3.1µm; GC-6.3µm. Morphological parameters like area, eccentricity (means how elongated the cell is, or how much the cell deviates from being a circle), irregularity (means how much the circumference of the cell deviates from the circumference of a perfect circle; 0= perfect circle, 1= highly irregular outline) have been decreased for cells grown on GC coated surface. While migration parameters, like-cell migration (means shortest distance between the starting point and the end point of the cell path), motility (means the actual distance traveled by the cell between the starting point and the end point) and motility speed (means motility distance divided by the time it took for the cell to travel from the starting point to the end point) has been increased for cells grown on GC coated surface. These changes are solely attributed to the glycation reaction that modified the native collagen. The changes in the cell growth pattern are also similar for HEK-RAGE cells grown on different coated surfaces. These two independent studies lead us to investigate the growth behavior of cancer cells on glycated ECM growth surface.

### Cancer cell morphology and migration analysis

For this experiment, we analyzed the growth behavior of four breast cancer cell lines-HCC1954 (data presented; for other cell lines see supplementary Fig. 8), Hs578T, HCC1143, and ZR-75-1 grown in three conditions: NT growth surface, NC coated growth surface, and GC coated growth surface. Following 24 hours of cell seeding, attached cells were monitored for another 24 hours. From our analysis, we found that on glycated growth surfaces, breast cancer cells behaved in a cell line-specific manner, though there are variations in cell morphology and migration among different growth surfaces.

HCC1954 is a poorly differentiated with HER2/neu+ overexpressed cell line that does not express estrogen or progesterone receptor. This cell line shows significant variations in cell growth among different growth surfaces (see Supplementary Fig. 6, online). Cells grown on the GC surface have decreased optical thickness and optical volume. The collagen glycation reduces average cell optical thickness and optical volume; for instance, the mean of optical thickness on NT-2.65 µm, NC-2.25 µm, and GC-2.33 µm, and the mean of optical volume on NT-2502 µm^3^, NC-2017 µm^3^, and GC-2113 µm^3^. A decrease in optical thickness corresponds with the increase in cell spreading area. Other morphological parameters such as cell spreading area increased on GC growth surface NT-975.5 µm^2^; NC-943.4 µm^2^; and GC-996.2 µm^2^, cell eccentricity decreased NT-0.635 µm; NC-0.630 µm; and GC-0.621 µm, cell irregularity increased NT-0.487 µm; NC-0.512 µm; and GC-0.525 µm and cell perimeter length increased NT-151.8 µm; NC-155.1 µm; and GC-160.8 µm (Fig. 8).

**Fig. 6.**
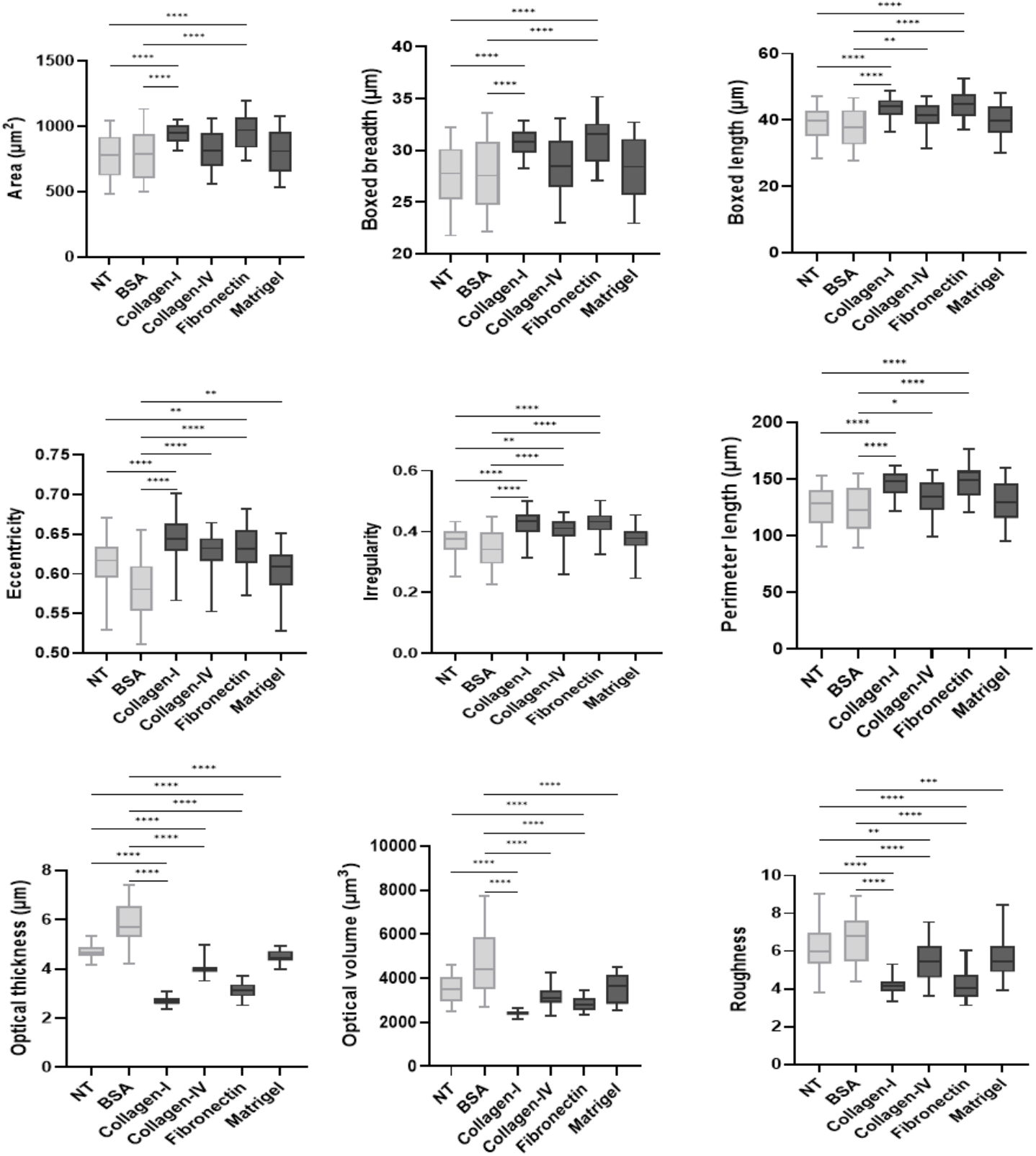
Differences in cell growth were observed in various ECM growth surfaces, with high significance. HEK293 cells grown in collagen I and fibronectin-coated growth surface show significant variations in almost all features compared to control groups. Data presented from two independent experiments. Adjusted p value > 0.05 (ns, non-significant, not mentioned in this figure), < 0.05 (*), < 0.01 (**), < 0.001 (***), or < 0.0001(****).

**Fig. 8.**
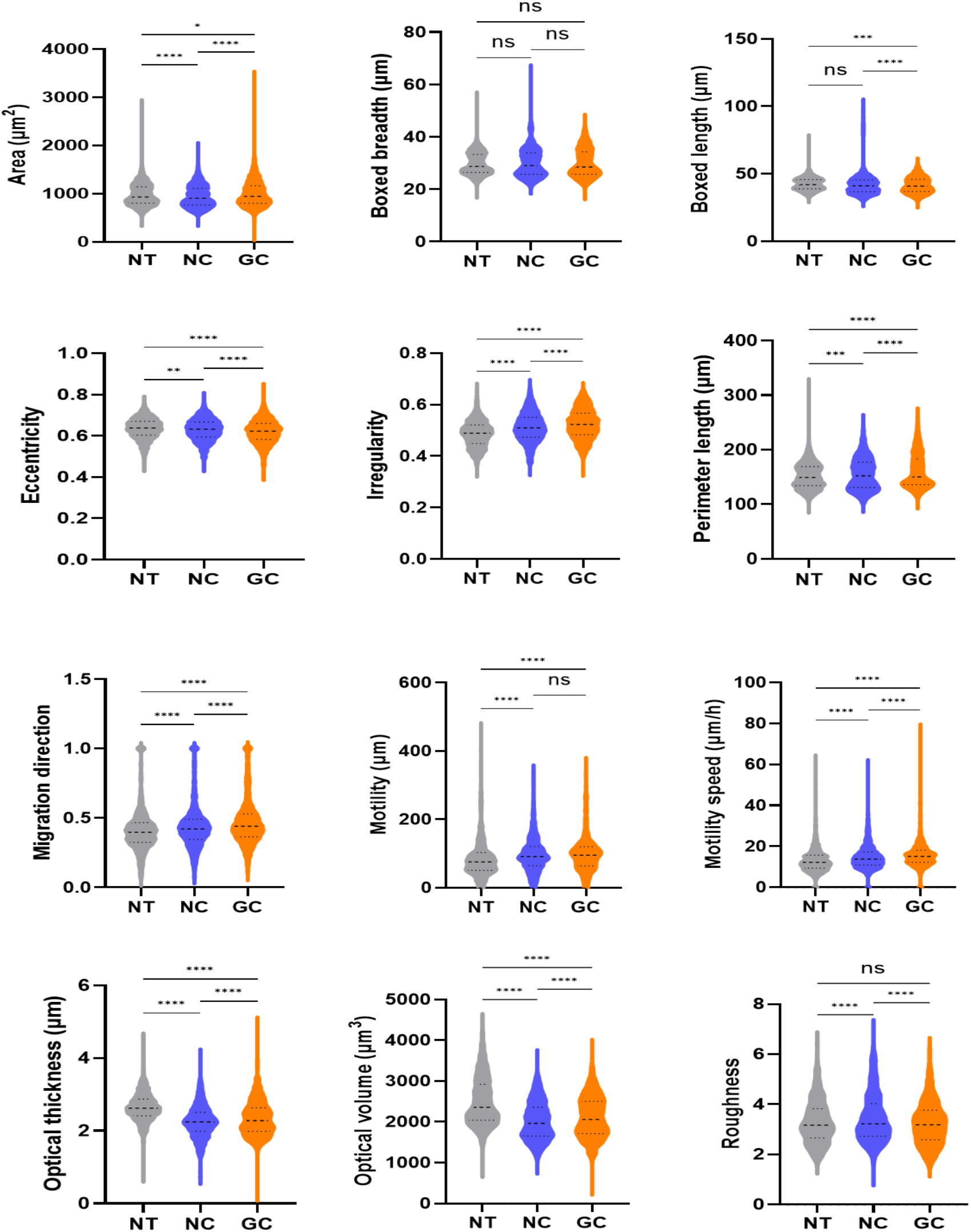
Holographic imaging of cells reveals the impact of ECM glycation on cell morphology and migration. Five randomly chosen positions were imaged at 20-minute intervals for 24 hours for each of the treatment conditions were analyzed. Differences in cancer cell morphology and migration resonate with co-related metastatic characteristics. Glycated growth surface-dependent changes in cell growth signify the importance of non-enzymatic glycation in cancer cell metastasis and invasion. Data presented from three independent experiments with replicates. Data presented mean ± SD from N=3 (with two replicates each) independent experiments. Adjusted p value > 0.05 (ns), < 0.05 (*), < 0.01 (**), < 0.001 (***), or < 0.0001(****).

To determine cells’ migratory parameters, the single-cell tracking feature of the AppSuite 3.5 software was utilized. In the case of migratory parameters, there were significant variations found in different growth surfaces. The average cell migration (Fig. 9d) increased on GC coated surface; mean of cell migration on NT-23.77 µm; NC-28.85 µm; and GC-30.4 µm, motility on NT-89.48 µm; NC-98.18 µm; and GC-99.83 µm and motility speed on NT-13.58 µm/h; NC-14.78 µm/h; GC-16.14 µm/h. Representative rose plot of single cell 2D movement trajectories from different growth surfaces, where cells are tracked for 24 hours (Fig. 9a, b, c).

**Fig. 9.**
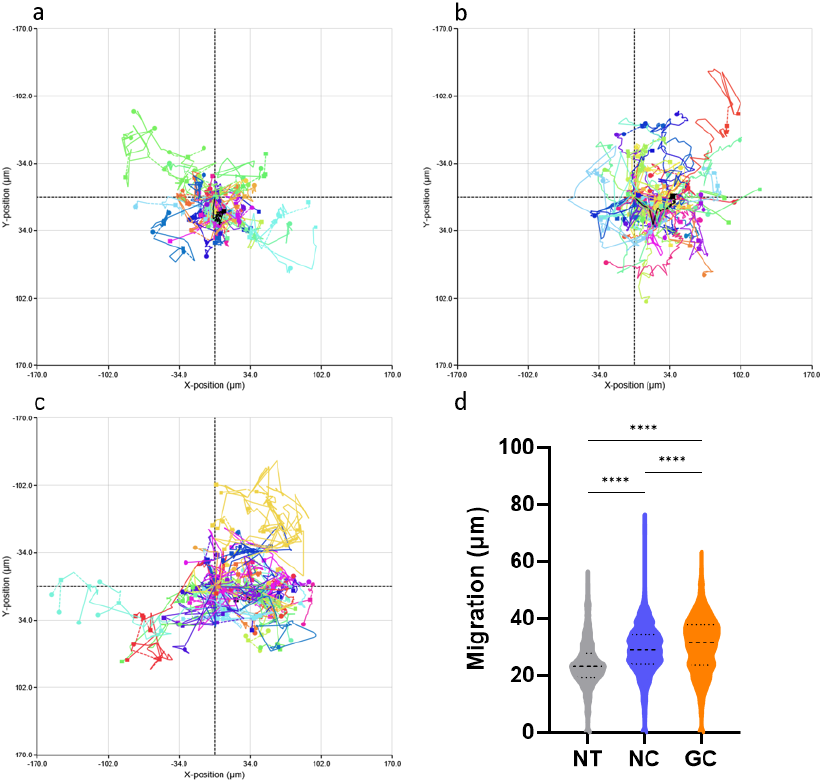
Rose plot representing the relative movement of HCC1954 cells on different growth surfaces. Fig. 9a, b, c represents NT, NC, and GC growth surfaces respectively. The relative single-cell movement plot for 24 hours is presented from one representative position of each condition. The cell’s movement plot corresponds with the average cells’ migration (Fig. 9. d) on different growth surfaces. The glycation of collagen significantly increased the migration of HCC1954 breast cancer cells.

Among the other cell lines, Hs578T is a non-tumorigenic breast cancer cell line with no estrogen expression and shows almost similar behavior to HCC1954 cells. HCC1143 is poorly differentiated with HER2/neu-cell line that does not express estrogen or progesterone receptors. This cell line has reduced adhesion on the GC growth surface. ZR-75-1 is a tumorigenic breast cancer cell line with estrogen expression. This cell line has a propensity to form clusters in the culture plate; as a result, some of the cell morphological features were not determined. Though in most cases, for all breast cancer cell lines, cells grown on the GC growth surface demonstrated some levels of variation compared to NC and NT conditions. Below (Fig. 10) is a representation of the optical thickness of all the cell lines on different growth surfaces.

**Fig. 10.**
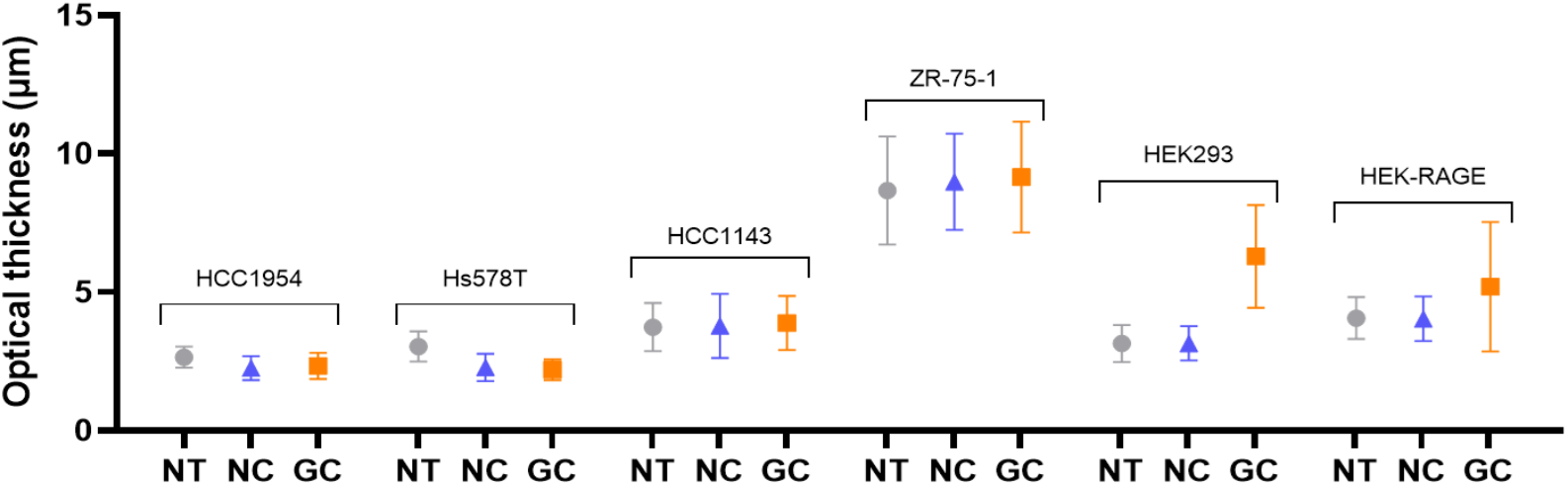
Changes in cell optical thickness on different growth surfaces for different cell lines.

**Fig. 11.**
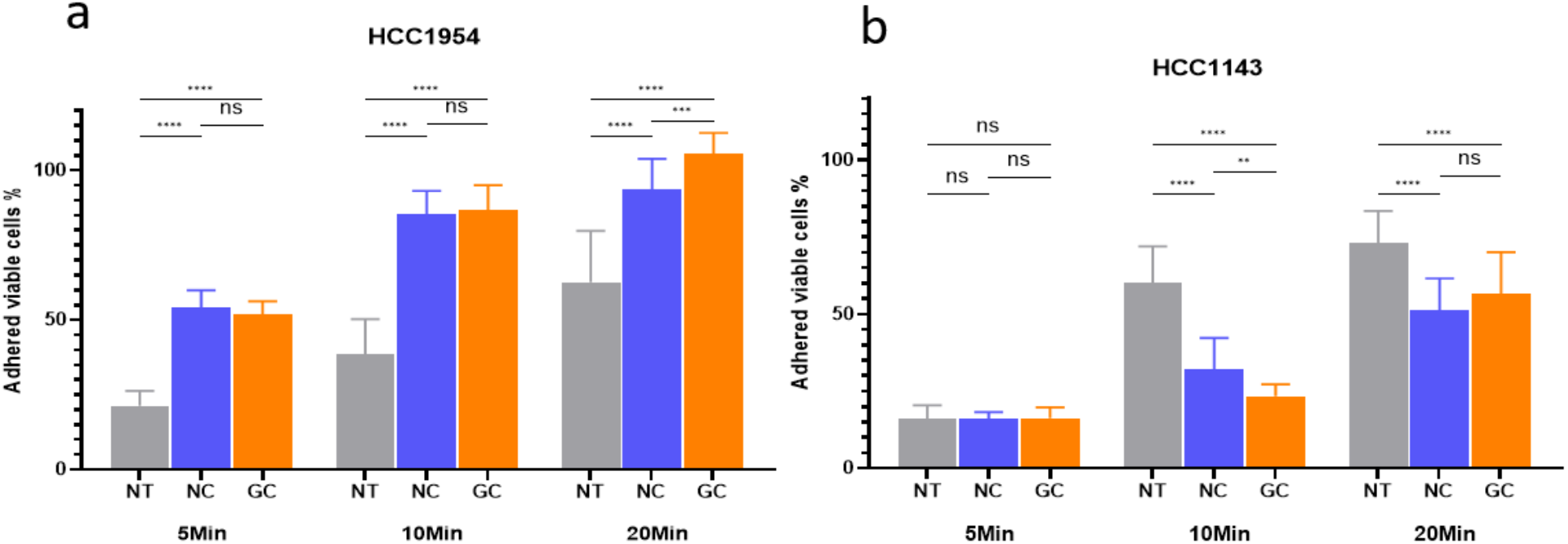
The plot of time-dependent cell adhesion of two breast cancer cells grown on NT, NC, and GC growth surfaces. Breast cancer cell adhesion with time significantly altered on the GC growth surface, and these changes are found to be cell line specific. Data presented mean fluorescence + SD of N=3 independent experiments. (For Hs578T and ZR-75-1 see Supplementary Fig. 7 online)

**Fig. 12.**
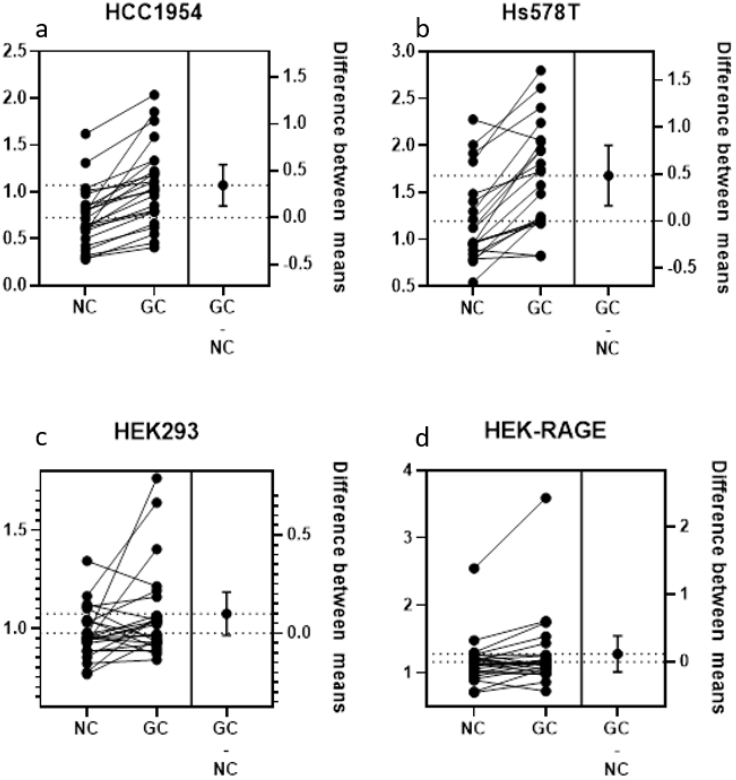
Estimation plot of relative fold change in gene expression for cells grown on NC and GC-coated growth surface.

### Effect of glycation on cell adhesion

To evaluate how the glycation of ECM affects the adhesion of cancer cells, we conducted a time-dependent cell adhesion assay, where breast cancer cells were incubated on NT, NC, and GC growth surfaces for 5 min, 10 min, and 20 min. After the incubation time, non-adherent cells were aspirated out, and the wells were washed twice with sterile PBS to ensure only the adhered cells remained in the wells. From the resazurin metabolic assay, we found that glycation of ECM alters the cancer cells’ adhesion properties. Such as for HCC1954 cells have higher adhesion for both NC and GC surfaces compared to NT surfaces, and with longer time GC surface shows higher cell adhesion properties. For HCC1143 cells, which is a low-adhesive cell line shown reduced cell adhesion in the NC and GC surfaces compared to the NT surface. These changes in cell adhesion properties are solely dependent on the attributed growth surfaces.

### Gene expression analysis of cells grown on different growth surfaces

To figure out whether the changes in the cell growth behavior are attributed to specific genes or it is just growth surface-dependent behavior changes, we analyzed the gene expression level of transcription and adhesion-related genes. For this, the experimental cell lines were grown on NT, NC, and GC-coated growth surfaces. After 48 hours of cell seeding, mRNA has been extracted and cDNA synthesized following standard procedures. A total of 29 gene expressions on different growth surfaces have been analyzed. After analyzing the relative fold change, we didn’t find any specific patterns in gene expression changes in cells grown on various growth surfaces. Though the breast cancer cells grew on the GC surface, for most genes, there was a slight increase in expression compared to NC. This makes the point clear that changes in cell morphology and migration are growth surface dependent, but not on other biological factors.

## Discussion

Here we used the DHM system that allowed continuous monitoring of cell growth in a cell culture incubator that can provide both qualitative and quantitative phase information on cells’ morphological and migration features. This system has an advantage over traditional techniques as it is a label-free non-invasive real-time imaging system as a result there is no need for the addition of dye or exogenous chemicals, and it permits cell monitoring in physiological and controlled conditions limiting the external environmental factors that may impact the results of the experiment and it allows to investigate cells behavior in single cell level^25,44^. In this study, we investigated the effect of MG-induced ECM glycation on cancer (breast) cell morphology and migration, highlighting the pivotal role of glycated ECM in modulating cancer progression. ECM is biologically long-lived and most vulnerable to glycation reactions and the generation of AGEs. The impact of ECM glycation has significant negative consequences in disease progression^9^. Our results revealed significant differences in cell morphological parameters between cells grown on NC vs GC. Holographic image analysis exhibits the changes in cell spreading area, eccentricity, irregularity, and perimeter length. These fundamental morphological parameter changes reflect the altered interaction between the ECM and cells^45^. Previous studies have demonstrated that glycation-mediated stiffening of ECM proteins modulates the interaction between ECM and growth factors, thereby altering cell adhesion and spreading^46-48^. From the single-cell migration analysis, we point out that cells grown on a glycated growth surface are comparably more motile than cells grown on an NC surface^49,50^. These findings are consistent with the established notion that glycated ECM promotes a pro-migratory environment, potentially due to the altered biochemical and mechanical cues presented by the glycated growth surface^51^. Interestingly, these changes in cell growth due to ECM glycation were found to be cell line-specific, suggesting that different cancer cell types may have distinct mechanisms for responding to changes in the ECM. This variability could stem from multiple interrelated factors such as growth factors and signaling molecules expression profiles, intracellular signaling pathways, or adaptive responses to ECM modifications. Literature studies show that the effect of ECM glycation on cells’ morphological behavior varies widely based on the cell types used, the experimental conditions employed, the types of ECM protein, and the glycation reagents used for the studies. Such as some studies reported that ECM glycation impaired the cell’s adhesion and migration capacity^6,47,52-54^ while others reported that glycation increased the cell adhesion strength, cell migration, and invasion^48,51,55-57^. Further studies are required to explore the molecular mechanism of these cell line-specific responses, specifically in the context of metastatic potential. These changes in cell morphology, adhesion, and migration, due to glycated ECM, may facilitate critical steps in tumor progression, including invasion and metastasis. Moreover, the pro-inflammatory milieu associated with glycated ECM may exacerbate cancer cell progression by promoting stromal activation and immune cell infiltration. These insights align with emerging evidence that AGEs and their receptors (RAGE) role in promoting cancer metastasis, invasion, and angiogenesis^58-60^.

While our study provides a comprehensive analysis of the effects of glycated ECM on cancer cell growth behavior, certain limitations need to be addressed in future investigations. First, we use a single ECM protein (collagen), which may not fully capture the complexity of the native tumor microenvironment, given the magnitude of ECM components and their dynamic interactions. Future studies should explore the effects of glycation on other ECM proteins, such as fibronectin and laminin, as well as the combined effects of multiple glycation agents. In addition, investigating the downstream signaling pathways influenced by glycated ECM could provide deeper mechanistic insights regarding cell line-specific behavior. The effect of glycation in in vivo conditions is required to confirm the relevance of our findings in a physiological and therapeutic context.

In conclusion, our study emphasizes the potential of using DHM for the evaluation of morphological and migration characteristics of cancerous cells. From the amino acid analysis, TEM imaging, and immunoreactivity assay, we showed the impact of MG glycation on ECM collagen. From the real-time holographic image analysis, we demonstrated that ECM glycation significantly impacts the cells’ physiological features on a single-cell level, which are directly correlated with cancer cell invasion and metastasis. Our cell adhesion and gene expression studies implied that the changes in cell behaviors are solely growth surface dependent, and this study will provide strong reasoning to incorporate the effect of ECM glycation in future cancer research and therapeutic studies.

## Materials and methods

### Collagen glycation and culture plate preparation

The fibrous collagen formation process involved the elevation of pH to 7.4, transitioning from the monomeric collagen (#L7213, Sigma-Aldrich) previously dissolved in an acidic solution. A coating of 10 μg/cm^2^ fibrous collagen was meticulously applied onto 6-well cell culture plates (#83.3920, Sarstedt). Subsequently, these plates were carefully air-dried on a laminar-flow hood (LFH). To glycate the coated collagen, 2 ml of 20 mM MG solution (#B24664, Alfa Aesar) was added into designated wells, while non-glycated wells received 2 ml of deionized (DI) water. The plates were carefully sealed using a plate sealer and placed in an incubator at 37°C for 3 days to facilitate the collagen glycation on the coated surfaces. Upon the completion of the glycation process, the plates were washed four times with DI water. Following the washing steps, the plates were air-dried and used immediately, or sealed with a plate sealer and stored at 4°C until subsequent use. During the process of cell seeding, an additional wash of the plates was carried out using the culture media.

For the identification of DHM-derived features, we used different ECM to coat the growth surface such as Collagen-I (50 μg/ml; #344702001, R&D Systems), Collagen-IV (5 μg/ml; # 3410-010-01, R&D Systems), Fibronectin (5 μg/ml; #3420-001-01, R&D Systems), Matrigel (10 μg/ml; #343300101, R&D Systems) with Bovine serum albumin (BSA) (100 μg/ml) and non-treated growth surface as control. After 24 hours of coating at room temperature inside a laminar flow hood (LFH), coating solutions were removed, the plate was washed once with water, and the plate was seeded with cells immediately.

### Amino acid analysis and transmission electron microscopy (TEM)

In preparation for amino acid analysis, after fibrous collagen formation (1 mg/ml), a 20 mM MG solution was introduced to the mixture and maintained within an incubator at a temperature of 37°C for 3 days. Following the glycation process, the solutions underwent a dialysis procedure against water, spanning a period of 2 days at a controlled temperature of 4°C. Throughout the dialysis, the water was periodically exchanged to ensure optimal purification. Upon completion of the dialysis, the contents were subjected to freeze-drying on a vacuum pump for a duration of 24 hours. Finally, the dried samples were dispatched for comprehensive amino acid analysis.

For TEM, both NC and GC samples were meticulously affixed onto specialized microscopic grids. Subsequently, the structural characteristics of these samples were thoroughly examined utilizing TEM techniques.

### Immunoassay

ELISA plates (#655061, GBO) were coated with collagen at a concentration of 10 μg/cm^2^ with a 24-hour air-drying process within an LFH. Glycation solution was carefully added into the wells, ranging from 20 mM to 10 μM in concentration, and the plates were kept at a temperature of 37°C for 3 days. Following glycation, the plates were washed 3 times and blocked with 3% BSA for 1 hour. The concentration gradient level of glycation was determined using 1-hour incubation with primary antibody (Mouse anti-methylglyoxal monoclonal antibody, #STA-011, Cell Biolabs) followed by 1-hour incubation with AP-conjugated secondary antibody (Alkaline Phosphatase AffiniPure Goat Anti-Mouse IgG, F(ab’)_2_ fragment specific, #115-055-072, Jackson ImmunoResearch). Finally, 1 mg/ml p-nitrophenol containing alkaline phosphatase buffer was used as substrate, and the plate was incubated at 37°C for color development. The absorbance was measured at a wavelength of 450nm using a spectrophotometer.

### General cell culture procedure

Breast cancer cell lines, HCC1954 (#CRL-2338, ATCC), HCC1143 (#CRL-2321, ATCC), and ZR-75-1 (#CRL-1500, ATCC), were cultured using RPMI (#30-2001, ATCC) and Hs578T (#HTB-126, ATCC), HEK293 (#CRL-1573, ATCC) and HEK-RAGE (stable RAGE expressing cells) were cultured using DMEM (#30-2002, ATCC) medium, both supplemented with 10% FBS (#PS-FB1, Peak serum), 1% Penicillin/Streptomycin (#K952, VWR) and for HEK-RAGE cells additional 0.05% G418 used. These cells were maintained within a humidified incubator at a temperature of 37°C, under a controlled 5% CO_2_ environment. To ensure optimal growth, the cells were passaged approximately every 4-5 days, with consistent monitoring of mycoplasma contamination via PCR. For the experimental procedures, the cells were seeded onto 6-well plates coated with GC at a density of 8×10^4^ cells per well, using 2 ml of growth media. After an initial 24 hours following cell seeding, the media were aspirated and subsequently replaced with 3 ml of fresh growth media. The traditional cell culture plate lid was replaced with transparent Hololid (Phase Holographic Imaging, Lund, Sweden) and plates were incubated for 30-45 min to eliminate condensation. The kinetic proliferation or morphology assay in AppSuite 3.5 (Phase Holographic Imaging, Lund, Sweden) was used to determine the cell morphology and migration, and the cells were monitored for a continuous 24-hour period.

### Quantitative phase imaging

Quantitative Phase Imaging was performed using a Holomonitor M4 microscope (Phase Holographic Imaging AB, Lund, Sweden). Images were segmented and analyzed using the AppSuite 3.5 software, and a total of 12 cell features covering cell morphology and migration were analyzed. Measured features include cell area, boxed length and breadth, eccentricity, irregularity, optical thickness, optical volume, cell migration, migration direction, cell motility, motility speed, perimeter length, and roughness. For cell segmentation, auto minimum error features were used for all experiments, and cell object size was determined based on individual cell lines and kept constant for representing cell lines.

### Cell adhesion assay

96-well Tissue culture plates were coated with 10 μg/cm^2^ fibrous collagen and air-dried on a laminar-flow hood (LFH). To glycate the coated collagen, 50 μL of 20 mM MG solution (#B24664, Alfa Aesar) was added into designated wells, while non-glycated wells received 50 μL of deionized (DI) water. The plates were carefully sealed using a plate sealer and placed in an incubator at 37°C for 3 days. Upon the completion of the glycation process, the plates were washed five times with DI water. Following the washing steps, the culture plates were seeded with 3×10^4^ cells in 100 μL of complete media. Immediately following seeding, cells were incubated at 37°C for these respective time points: 5, 10, and 20 min. After the incubation time, the cell suspension from designated wells was aspirated out and washed twice with sterile PBS. After that, wells were filled with 100 μL of complete media, and 10% V/V of resazurin (1 mg/ml) solution was added after the final wash and incubated at 37°C for 4 hours. The resazurin fluorescence was measured at 540/590 nm (excitation/emission) using a SpectraMax multi-plate reader, and the percentage of adhered cells was calculated using the formula-% Cells adhered = (Fluorescence from washed well/non-washed well) x 100

### qPCR assay

Cells were seeded on a GC-coated 6-well plate (#10062-892, VWR), at a density of 1.5×10^5^ cells per well with 2 mL of growth medium. After 48 hours of cell seeding media were aspirated, the cell monolayer was washed with 1x PBS, and the RNA was extracted using (#R6834-02, Omega E.Z.N.A.® Total RNA Kit II). Following RNA extraction, the RNAs (1 µg) were reverse transcribed into cDNA using reverse transcription reagents from NEB (#M0253, NEB) and stored at -20 °C. qPCR was performed using 20 ng of cDNA on a Stratagene Mx3000p thermocycler using the 5x HOT FIREPol EvaGreen qPCR Mix Plus (ROX) (#08-24-00001, Solis Biodyne) using appropriate primers. The S18 gene was used as the reference gene for fold change calculation. The following qPCR program was used: 12 min at 95 °C followed by 35 cycles of 15 sec at 95 °C, 20 sec at 62 °C, 20 sec at 72 °C. A melting curve was recorded at the end of the cycles to evaluate the quality of the amplified products. The fold (F) of change in gene expression was calculated for each gene using the ΔΔCt method with F = 2^(ΔΔCt).^ Where ΔCt = Ct _Gene_ – Ct _S18_, ΔΔCt= ΔCt_Treatment_-ΔCt_Control._

### Statistical analysis

Statistical analysis was performed using GraphPad Prism version 10.4.0 for Windows (San Diego, CA, USA). One-way ANOVA with Bonferroni’s multiple comparisons test was used to determine significant differences in unique feature means between the control and the treatments. Results are expressed as mean ± standard deviation unless mentioned otherwise. Adjusted p value > 0.05 (ns; non-significant), < 0.05 (*), < 0.01 (**), < 0.001 (***), or < 0.0001(****).

## Data availability

All processed data generated or analyzed during this study are included in this published article and its supplementary information files. Additional raw data from measurements are available upon request from the corresponding author.

## Author contributions

Conceptualization, Study Design, Methodology, Data Collection, Manuscript Writing, Experimental Implementation, Manuscript Editing and Manuscript Revision.

All authors approved the manuscript for publication.

## Declarations

## Additional information

Supplementary Information

## References

1 Theocharis, A. D., Skandalis, S. S., Gialeli, C. & Karamanos, N. K. Extracellular matrix structure. Advanced Drug Delivery Reviews 97, 4–27 (2016). 10.1016/j.addr.2015.11.001

2 Geiger, B. & Yamada, K. M. Molecular Architecture and Function of Matrix Adhesions. Cold Spring Harbor Perspectives in Biology 3 (2011). 10.1101/cshperspect.a005033

3 Yamada, K. M. Extracellular matrix dynamics in cell migration, invasion, and tissue morphogenesis. International Journal of Experimental Pathology 100, A4–A4 (2019).

4 Duran-Jimenez, B. et al. Advanced glycation end products in extracellular matrix proteins contribute to the failure of sensory nerve regeneration in diabetes. Diabetes 58, 2893–2903 (2009). 10.2337/db09-0320

5 Rabbani, N. & Thornalley, P. J. Glycation research in amino acids: a place to call home. Amino Acids 42, 1087–1096 (2012). 10.1007/s00726-010-0782-1

6 Haucke, E., Navarrete-Santos, A., Simm, A., Silber, R. & Hofmann, B. Glycation of extracellular matrix proteins of immune cells. Wound Repair and Regeneration 22, 239–245 (2014). 10.1111/wrr.12144

7 Yuan, Z. et al. Extracellular matrix remodeling in tumor progression and immune escape: from mechanisms to treatments. Mol Cancer 22, 48 (2023). 10.1186/s12943-023-01744-8

8 Naba, A., Clauser, K. R., Lamar, J. M., Carr, S. A. & Hynes, R. O. Extracellular matrix signatures of human mammary carcinoma identify novel metastasis promoters. Elife 3 (2014). 10.7554/eLife.01308

9 Rojas, A., Anazco, C., Gonzalez, I. & Araya, P. Extracellular matrix glycation and receptor for advanced glycation end-products activation: a missing piece in the puzzle of the association between diabetes and cancer. Carcinogenesis 39, 515–521 (2018). 10.1093/carcin/bgy012

10 Acerbi, I. et al. Human breast cancer invasion and aggression correlates with ECM stiffening and immune cell infiltration. Integrative Biology 7, 1120–1134 (2015). 10.1039/c5ib00040h

11 Leight, J. L., Wozniak, M. A., Chen, S., Lynch, M. L. & Chen, C. S. Matrix rigidity regulates a switch between TGF-beta 1-induced apoptosis and epithelial-mesenchymal transition. Molecular Biology of the Cell 23, 781–791 (2012). 10.1091/mbc.E11-06-0537

12 Bordeleau, F. et al. Matrix stiffening promotes a tumor vasculature phenotype. Proceedings of the National Academy of Sciences of the United States of America 114, 492–497 (2017). 10.1073/pnas.1613855114

13 Cross, K. et al. Role of the Receptor for Advanced Glycation End Products (RAGE) and Its Ligands in Inflammatory Responses. Biomolecules 14 (2024).

14 Schroter, D. & Hohn, A. Role of Advanced Glycation End Products in Carcinogenesis and their Therapeutic Implications. Current Pharmaceutical Design 24, 5245–5251 (2018). 10.2174/1381612825666190130145549

15 Eva, T. et al. Perspectives on signaling for biological- and processed food-related advanced glycation end-products and its role in cancer progression. Critical Reviews in Food Science and Nutrition 62, 2655–2672 (2022). 10.1080/10408398.2020.1856771

16 Vetter, S. W. Glycated Serum Albumin and AGE Receptors. Advances in Clinical Chemistry 72, 205–275 (2015). 10.1016/bs.acc.2015.07.005

17 Reddy, V., Aryal, P. & Darkwah, E. Advanced Glycation End Products in Health and Disease. Microorganisms 10 (2022). 10.3390/microorganisms10091848

18 Prasad, C., Davis, K., Imrhan, V., Juma, S. & Vijayagopal, P. Advanced Glycation End Products and Risks for Chronic Diseases: Intervening Through Lifestyle Modification. American Journal of Lifestyle Medicine 13, 384–404 (2019). 10.1177/1559827617708991

19 Yan, S., Ramasamy, R. & Schmidt, A. Mechanisms of disease: advanced glycation end-products and their receptor in inflammation and diabetes complications. Nature Clinical Practice Endocrinology & Metabolism 4, 285–293 (2008). 10.1038/ncpendmet0786

20 Twarda-Clapa, A., Olczak, A., Bialkowska, A. & Koziolkiewicz, M. Advanced Glycation End-Products (AGEs): Formation, Chemistry, Classification, Receptors, and Diseases Related to AGEs. Cells 11 (2022). 10.3390/cells11081312

21 Ricard-Blum, S. The Collagen Family. Cold Spring Harbor Perspectives in Biology 3 (2011). 10.1101/cshperspect.a004978

22 Sun, B. The mechanics of fibrillar collagen extracellular matrix. Cell Reports Physical Science 2 (2021). 10.1016/j.xcrp.2021.100515

23 Hamanaka, R. B. & Mutlu, G. M. The role of metabolic reprogramming and de novo amino acid synthesis in collagen protein production by myofibroblasts: implications for organ fibrosis and cancer. Amino Acids 53, 1851–1862 (2021). 10.1007/s00726-021-02996-8

24 Nash, A., Noh, S. Y., Birch, H. L. & de Leeuw, N. H. Lysine-arginine advanced glycation end-product cross-links and the effect on collagen structure: A molecular dynamics study. Proteins-Structure Function and Bioinformatics 89, 521–530 (2021). 10.1002/prot.26036

25 Hellesvik, M., Oye, H. & Aksnes, H. Exploiting the potential of commercial digital holographic microscopy by combining it with 3D matrix cell culture assays. Scientific Reports 10 (2020). 10.1038/s41598-020-71538-1

26 Huang, D. A. et al. Identifying fates of cancer cells exposed to mitotic inhibitors by quantitative phase imaging. Analyst 145, 97–106 (2020). 10.1039/c9an01346f

27 Hejna, M., Jorapur, A., Song, J. & Judson, R. High accuracy label-free classification of single-cell kinetic states from holographic cytometry of human melanoma cells. Scientific Reports 7 (2017). 10.1038/s41598-017-12165-1

28 Barker, K., Boucher, K. & Judson-Torres, R. Label-Free Classification of Apoptosis, Ferroptosis and Necroptosis Using Digital Holographic Cytometry. Applied Sciences 10 (2020). 10.3390/app10134439

29 Chong, S. et al. Methylglyoxal inhibits the binding step of collagen phagocytosis. Journal of Biological Chemistry 282, 8510–8520 (2007). 10.1074/jbc.M609859200

30 Yuen, A. et al. Methylglyoxal-modified collagen promotes myofibroblast differentiation. Matrix Biology 29, 537–548 (2010). 10.1016/j.matbio.2010.04.004

31 Nagai, R. et al. Glycolaldehyde, a reactive intermediate for advanced glycation end products, plays an important role in the generation of an active ligand for the macrophage scavenger receptor. Diabetes 49, 1714–1723 (2000). 10.2337/diabetes.49.10.1714

32 Thorpe, S. & Baynes, J. Maillard reaction products in tissue proteins: New products and new perspectives. Amino Acids 25, 275–281 (2003). 10.1007/s00726-003-0017-9

33 Oya, T. et al. Methylglyoxal modification of protein - Chemical and immunochemical characterization of methylglyoxal-arginine adducts. Journal of Biological Chemistry 274, 18492–18502 (1999).

34 Morgan, P., Sheahan, P., Pattison, D. & Davies, M. Methylglyoxal-induced modification of arginine residues decreases the activity of NADPH-generating enzymes. Free Radical Biology and Medicine 61, 229–242 (2013). 10.1016/j.freeradbiomed.2013.03.025

35 Bose, T., Bhattacherjee, A., Banerjee, S. & Chakraborti, A. Methylglyoxal-induced modifications of hemoglobin: Structural and functional characteristics. Archives of Biochemistry and Biophysics 529, 99–104 (2013). 10.1016/j.abb.2012.12.001

36 Gao, Y. & Wang, Y. Site-selective modifications of arginine residues in human hemoglobin induced by methylglyoxal. Biochemistry 45, 15654–15660 (2006). 10.1021/bi061410o

37 Shinohara, M. et al. Overexpression of glyoxalase-I in bovine endothelial cells inhibits intracellular advanced glycation endproduct formation and prevents hyperglycemia-induced increases in macromolecular endocytosis. Journal of Clinical Investigation 101, 1142–1147 (1998).

38 Thornalley, P. J. Pharmacology of methylglyoxal: formation, modification of proteins and nucleic acids, and enzymatic detoxification-A role in pathogenesis and antiproliferative chemotherapy. General Pharmacology: The Vascular System 27, 565–573 (1996). /10.1016/0306-3623(95)02054-3

39 Kilhovd, B. et al. Increased serum levels of methylglyoxal-derived hydroimidazolone-AGE are associated with increased cardiovascular disease mortality in nondiabetic women. Atherosclerosis 205, 590–594 (2009). 10.1016/j.atherosclerosis.2008.12.041

40 Bansode, S. et al. Glycation changes molecular organization and charge distribution in type I collagen fibrils. Scientific Reports 10 (2020). 10.1038/s41598-020-60250-9

41 Fessel, G. et al. Advanced glycation end-products reduce collagen molecular sliding to affect collagen fibril damage mechanisms but not stiffness. PLoS One 9, e110948 (2014). 10.1371/journal.pone.0110948

42 Li, Y., Fessel, G., Georgiadis, M. & Snedeker, J. Advanced glycation end-products diminish tendon collagen fiber sliding. Matrix Biology 32, 169–177 (2013). 10.1016/j.matbio.2013.01.003

43 Schalkwijk, C., Micali, L. & Wouters, K. Advanced glycation endproducts in diabetes-related macrovascular complications: focus on methylglyoxal. Trends in Endocrinology & Metabolism 34, 49–60 (2023). 10.1016/j.tem.2022.11.004

44 Janicke, B., Kårsnäs, A., Egelberg, P. & Alm, K. Label-Free High Temporal Resolution Assessment of Cell Proliferation Using Digital Holographic Microscopy. Cytometry Part A 91A, 460–469 (2017). 10.1002/cyto.a.23108

45 Sloseris, D. & Forde, N. R. AGEing of collagen: The effects of glycation on collagen’s stability, mechanics and assembly. Matrix Biology 135, 153–160 (2025).

46 Wang, W. et al. Diabetic hyperglycemia promotes primary tumor progression through glycation-induced tumor extracellular matrix stiffening. Science Advances 8 (2022). 10.1126/sciadv.abo1673

47 McCarthy, A., Uemura, T., Etcheverry, S. & Cortizo, A. Advanced glycation endproducts interfere with integrin-mediated osteoblastic attachment to a type-I collagen matrix. International Journal of Biochemistry & Cell Biology 36, 840–848 (2004). 10.1016/j.biocel.2003.09.006

48 Mason, B. N., Starchenko, A., Williams, R. M., Bonassar, L. J. & Reinhart-King, C. A. Tuning three-dimensional collagen matrix stiffness independently of collagen concentration modulates endothelial cell behavior. Acta Biomater 9, 4635–4644 (2013). 10.1016/j.actbio.2012.08.007

49 Sharaf, H. et al. Advanced glycation endproducts increase proliferation, migration and invasion of the breast cancer cell line MDA-MB-231. Biochimica Et Biophysica Acta-Molecular Basis of Disease 1852, 429–441 (2015). 10.1016/j.bbadis.2014.12.009

50 Schildhauer, P. et al. Glycation Leads to Increased Invasion of Glioblastoma Cells. Cells 12 (2023). 10.3390/cells12091219

51 Koorman, T. et al. Spatial collagen stiffening promotes collective breast cancer cell invasion by reinforcing extracellular matrix alignment. Oncogene 41, 2458–2469 (2022). 10.1038/s41388-022-02258-1

52 Bartling, B., Desole, M., Rohrbach, S., Silber, R. & Simm, A. Age-associated changes of extracellular matrix collagen impair lung cancer cell migration. FASEB Journal 23, 1510–1520 (2009). 10.1096/fj.08-122648

53 Reigle, K. et al. Non-enzymatic glycation of type I collagen diminishes collagen-proteoglycan binding and weakens cell adhesion. Journal of Cellular Biochemistry 104, 1684–1698 (2008). 10.1002/jcb.21735

54 Said, G. et al. Impact of carbamylation and glycation of collagen type I on migration of HT1080 human fibrosarcoma cells. International Journal of Oncology 40, 1797–1804 (2012). 10.3892/ijo.2012.1393

55 Kemeny, S., Cicalese, S., Figueroa, D. & Clyne, A. Glycated collagen and altered glucose increase endothelial cell adhesion strength. Journal of Cellular Physiology 228, 1727–1736 (2013). 10.1002/jcp.24313

56 Kyurkchiev, S., Ivanov, G. & Manolova, V. Advanced glycosylated end products activate the functions of cell adhesion molecules on lymphoid cells. Cellular and Molecular Life Sciences 53, 911–916 (1997).

57 Suh, Y. et al. Glycation of collagen matrices promotes breast tumor cell invasion. Integrative Biology 11, 109–117 (2019). 10.1093/intbio/zyz011

58 Palanissami, G. & Paul, S. RAGE and Its Ligands: Molecular Interplay Between Glycation, Inflammation, and Hallmarks of Cancer-a Review. Hormones & cancer 9, 295–325 (2018). 10.1007/s12672-018-0342-9

59 Ahmad, S. et al. AGEs, RAGEs and s-RAGE; friend or foe for cancer. Seminars in Cancer Biology 49, 44–55 (2018). 10.1016/j.semcancer.2017.07.001

60 Logsdon, C., Fuentes, M., Huang, E. & Arumugam, T. RAGE and RAGE ligands in cancer. Current Molecular Medicine 7, 777–789 (2007).

