## Supplemental Information for "Using digital holographic microscopy (DHM) to monitor effects of extracellular matrix (ECM) glycation on cancer cell morphology and migration"

**Supplementary Fig. 1.** Glycated collagen (GC) TEM images.

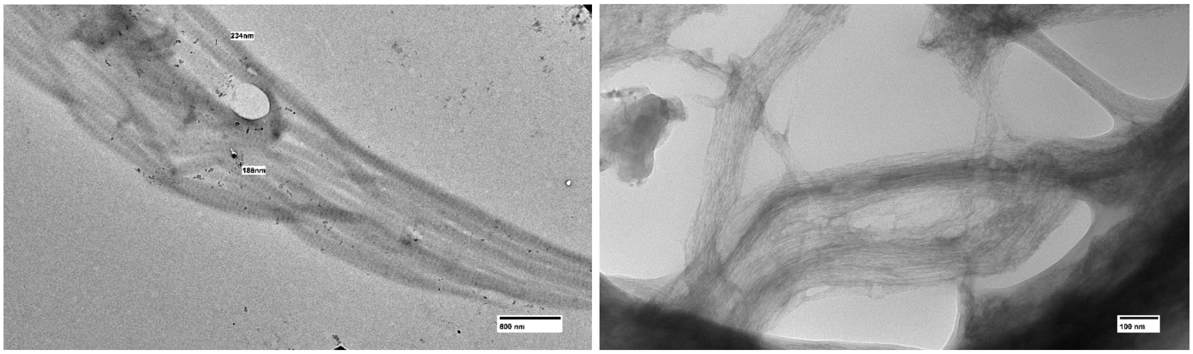


**Supplementary Fig. 2.** HEK293 cells on different growth surfaces- BSA, Collagen-IV, Fibronectin, and Matrigel, respectively.

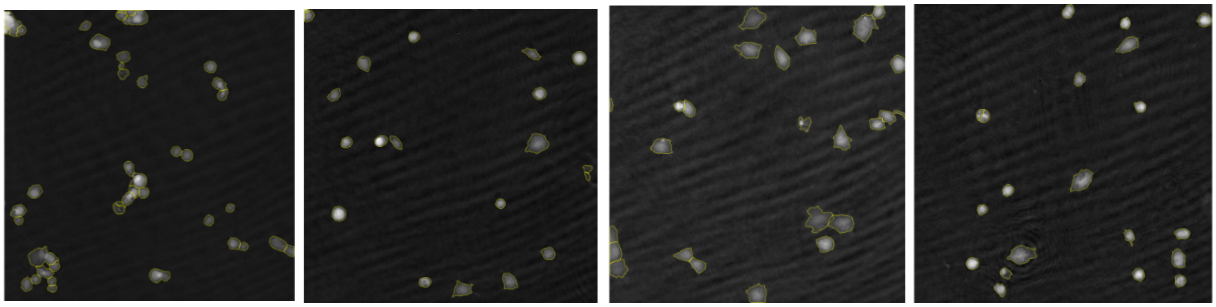

**Supplementary Fig. 3.** Differences in cell growth were observed in various ECM growth surfaces, with high significance. HEK-RAGE cells grown in collagen I and fibronectin-coated growth surface show significant variations in almost all features compared to control groups. Data presented from three independent experiments. Adjusted p value > 0.05 (ns, non-significant, not mentioned in this figure), < 0.05 (*), < 0.01 (**), < 0.001 (***), or < 0.0001(****).

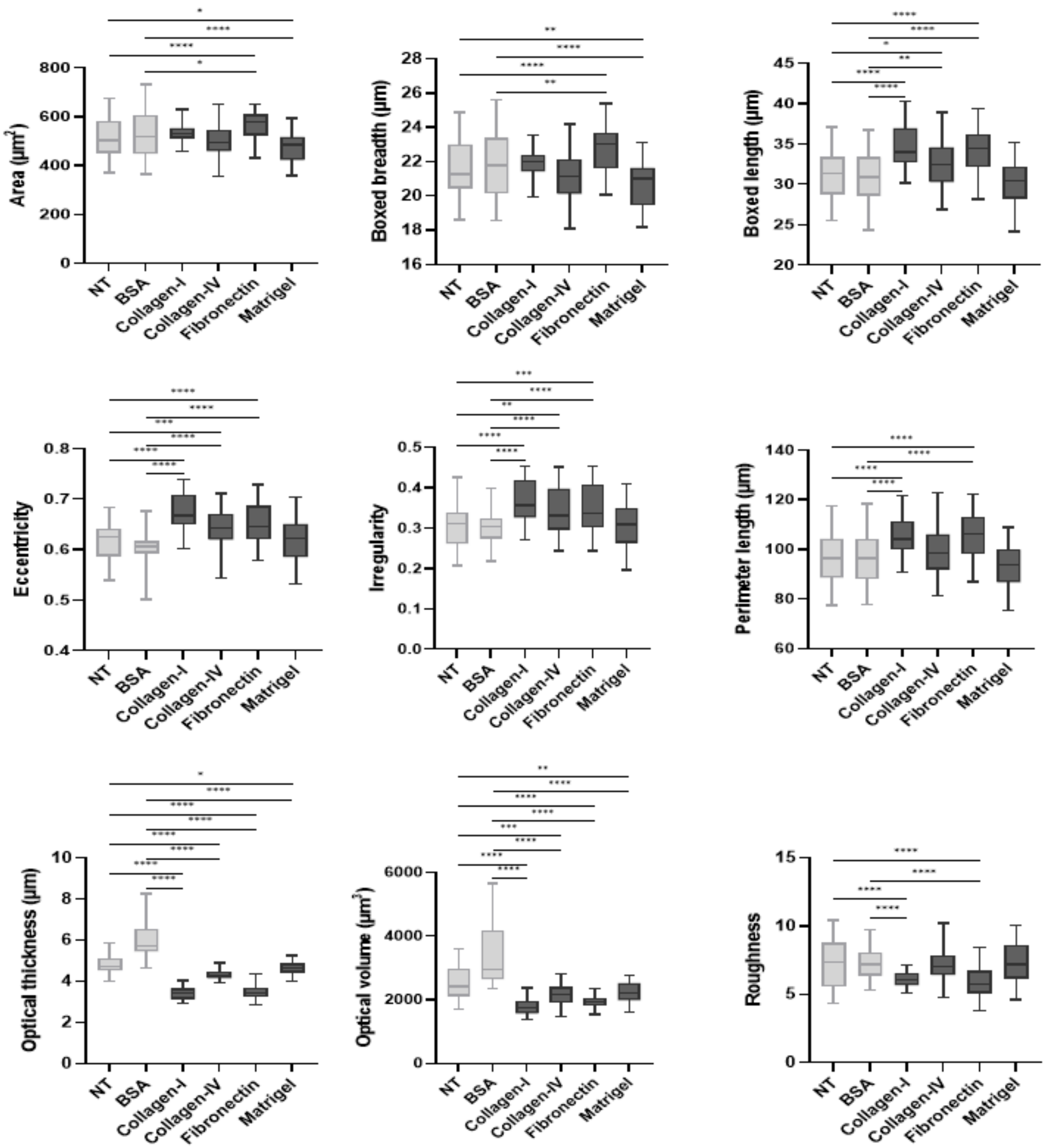

**Supplementary Fig. 4.** HEK293 cells grown on Non-treated (NT), non-glycated collagen (NC) vs glycated collagen (GC) coated growth surfaces.

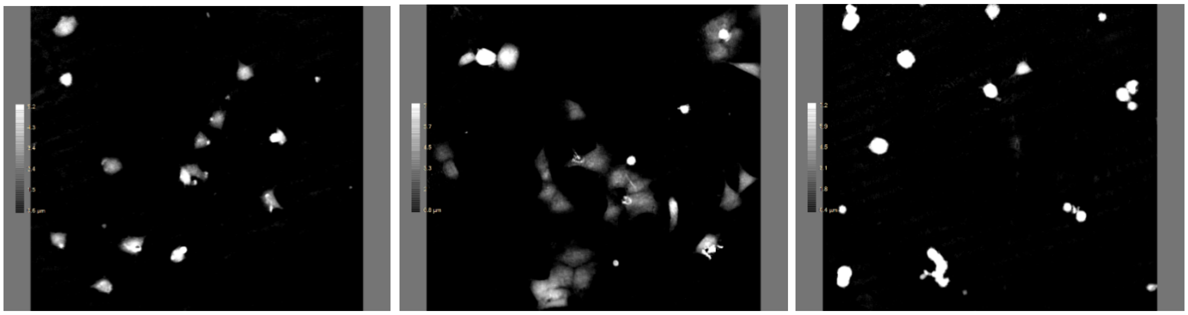

Representative HEK293 single cell from the GC growth surface with high optical thickness and optical volume.

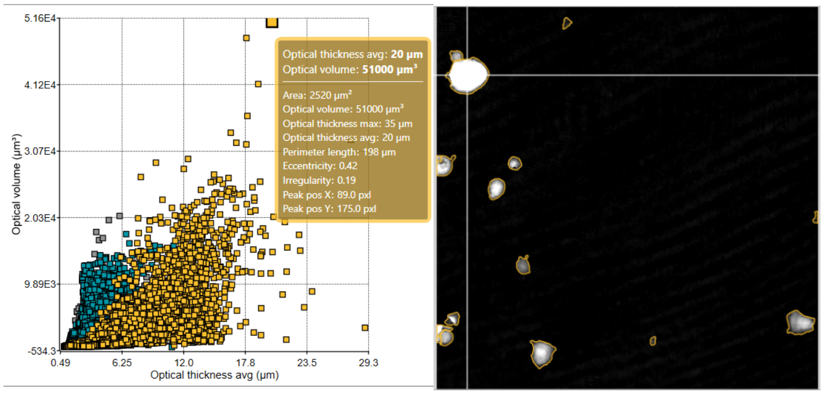

**Supplementary Fig. 5.** Holographic imaging of HEK293 cells reveals the impact of ECM glycation on cell morphology and migration. Five randomly chosen positions were imaged at 20-minute intervals for 24 hours for each of the treatment conditions were analyzed. Differences in cancer cell morphology and migration resonate with co-related metastatic characteristics. Glycated growth surface-dependent changes in cell growth signify the importance of non-enzymatic glycation in cancer cell metastasis and invasion. Data presented from three independent experiments with replicates. Data presented mean ± SD from N=3 (with two replicates each) independent experiments. Adjusted p value > 0.05 (ns), < 0.05 (*), < 0.01 (**), < 0.001 (***), or < 0.0001(****).

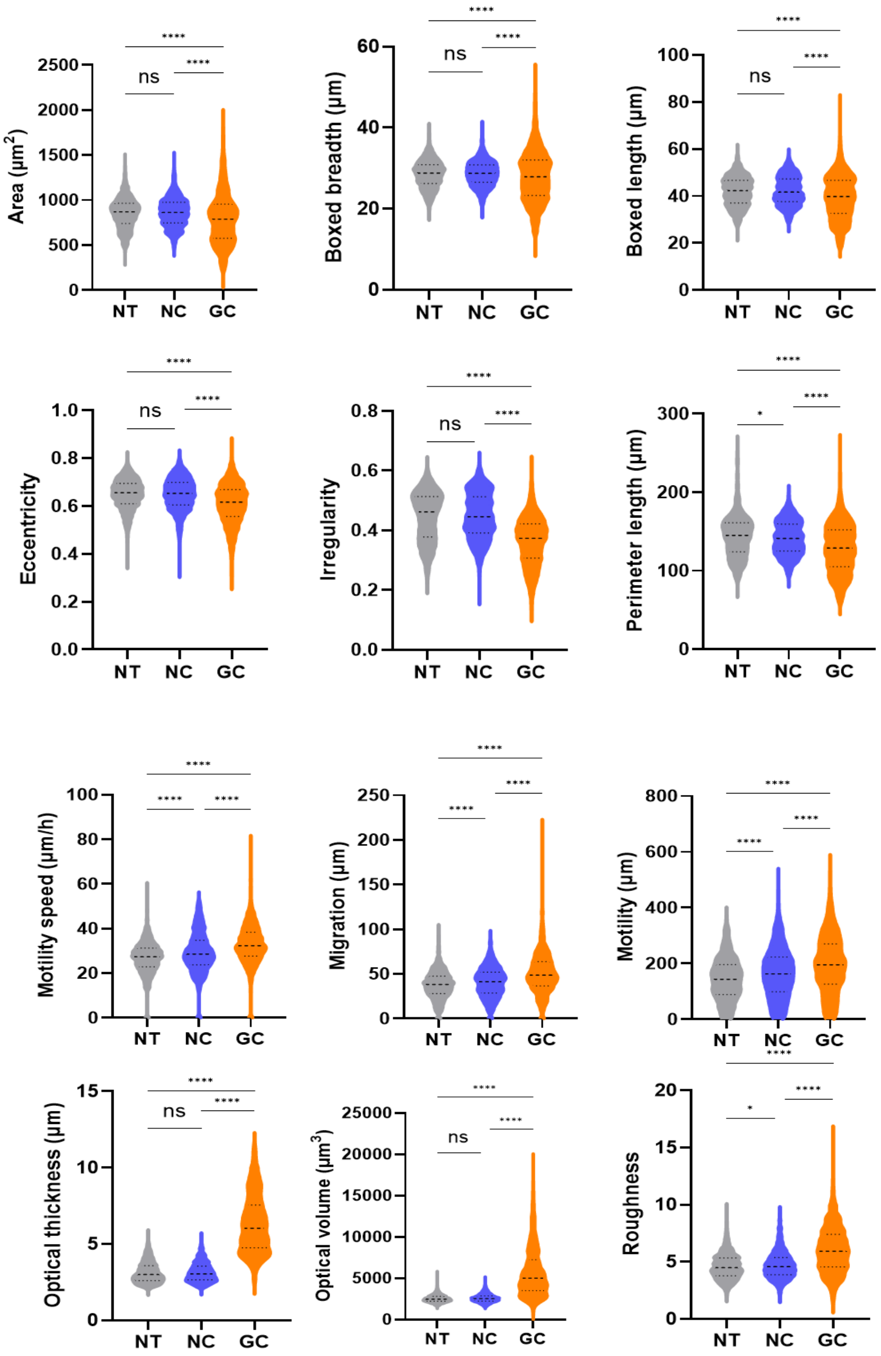

**Supplementary Fig. 6.** HCC1954 cells grown on Non-treated (NT), non-glycated collagen (NC) vs glycated collagen (GC) coated growth surfaces.

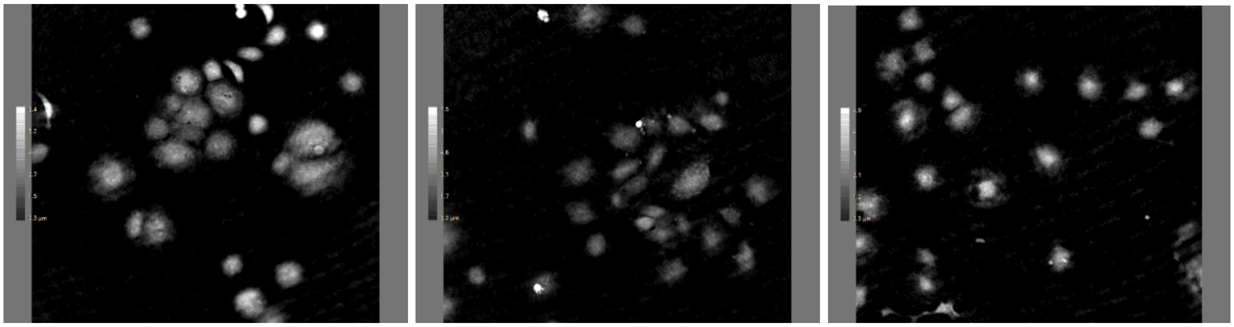

Representative single cell from NT growth surface with high optical thickness and optical volume.

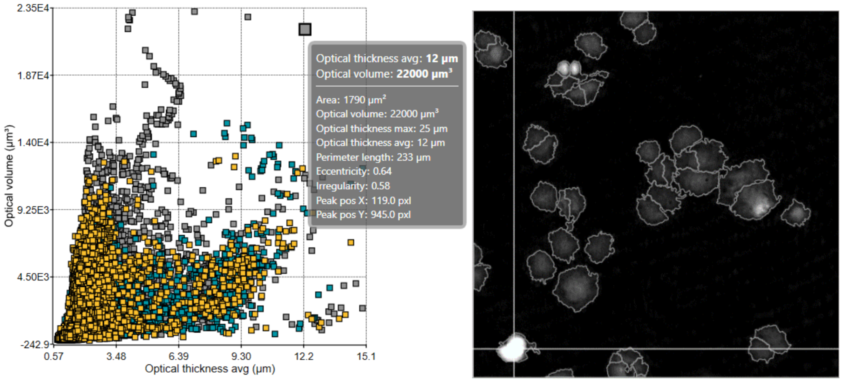

**Supplementary Fig. 7.** The time-dependent cell adhesion of breast cancer cells grown on NT, NC, and GC growth surfaces. Breast cancer cell adhesion with time significantly altered on the GC growth surface, and these changes are found to be cell line specific. Data presented mean fluorescence + SD of N=3 independent experiments.

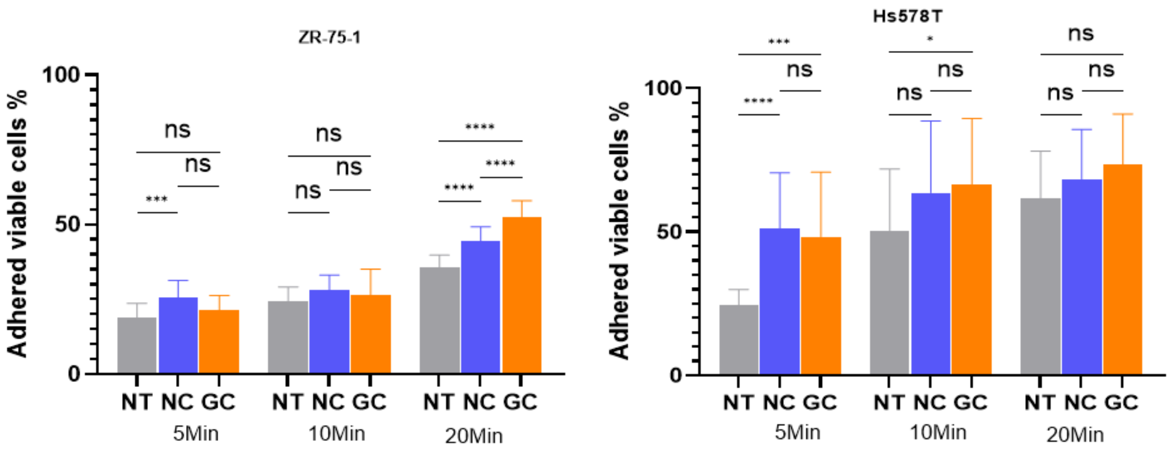

**Supplementary Fig. 8.** Among other cell lines, Hs578T and HCC1143 are breast cancer cell lines, and HEK-RAGE is an RAGE overexpressing cell line. Changes in cell growth behavior on different growth surfaces.

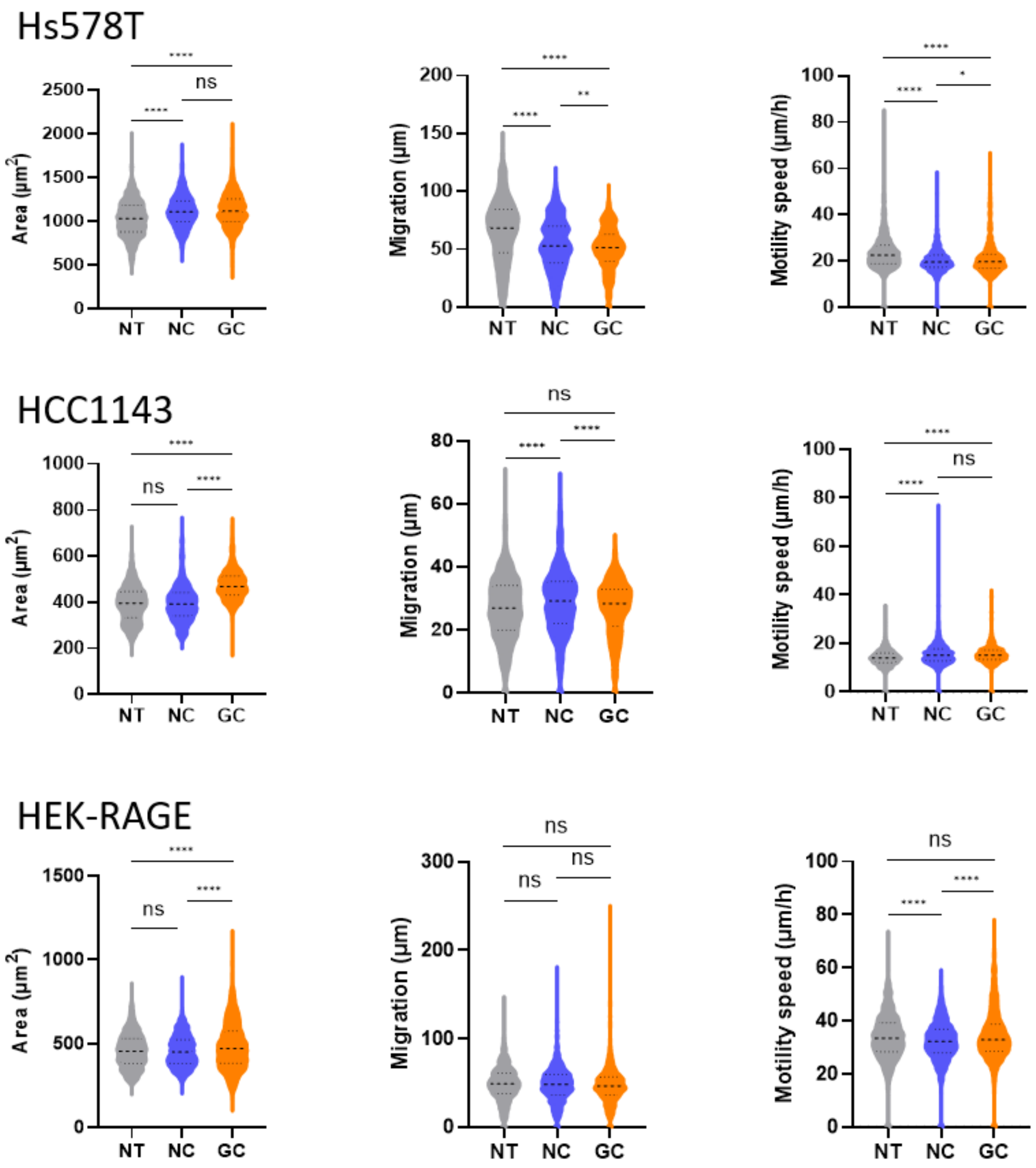
